# Intestinal epithelial IPMK protects mice from experimental colitis via governing colonic tuft cell development

**DOI:** 10.1101/2021.09.15.460418

**Authors:** Seung Eun Park, Jae Woong Jeong, Su-Hyung Lee, Seung Ju Park, Jaeseung Ryu, Se Kyu Oh, Sungsoon Fang, Seyun Kim

**Affiliations:** Department of Biological Sciences, Korea Advanced Institute of Science and Technology (KAIST), Daejeon, Republic of Korea; Department of Medicine, Yonsei University College of Medicine, Seoul, Republic of Korea; Section of Surgical Sciences, Vanderbilt University Medical Center, Nashville, TN 37232-2733, USA; Epithelial Biology Center, Vanderbilt University Medical Center, Nashville, TN 37232-2733, USA; KYNOGEN Co., Suwon, Republic of Korea; Severance Biomedical Science Institute, Graduate School of Medical Science, Brain Korea 21 Project for Medical Science, Yonsei University College of Medicine, Seoul, Republic of Korea; KAIST Institute for the BioCentury, KAIST, Daejeon, Republic of Korea

**Keywords:** Colon, IPMK, Epithelial barrier, Tuft cell, Colitis, IBD

## Abstract

As a pleiotropic signaling factor, inositol polyphosphate multikinase (IPMK) is involved in key biological events such as growth and innate immunity, acting either enzymatically to mediate the biosynthesis of inositol polyphosphates and phosphatidylinositol 3,4,5-trisphosphates, or noncatalytically to control key signaling target molecules. However, the functional significance of IPMK in regulating gut epithelial homeostasis remains largely unknown. Here we show that intestinal epithelial-specific deletion of IPMK aggravates dextran sulfate sodium (DSS)-induced colitis with higher clinical colitis scores and elevated epithelial barrier permeability. No apparent defects in PI3K-AKT signaling pathway and pro-inflammatory cytokine production were found in IPMK-deficient colons challenged by DSS treatment. RNA-sequencing and FACS analyses further revealed significantly decreased tuft cells in IPMK-deficient colons. Importantly, IPMK deletion in the gut epithelium was found to decrease choline acetyltransferase (ChAT) but not IL-25, suggesting selective loss of cholinergic signaling. Thus, these findings identify IPMK as a physiological determinant of tuft cell differentiation and highlight the critical function of IPMK in the control of gut homeostasis.

## Introduction

The gastrointestinal (GI) tract is lined by a constantly changing population of intestinal epithelial cells (IECs). Mammalian IECs exhibit a multitude of physiological functions, including absorption of nutrients, secretion of hormones as well as neurotransmitters, production of antimicrobial peptides in response to infectious stimuli, and control of tissue regeneration after physical damage. The intestinal mucosa undergoes a process of continual renewal characterized by active proliferation of crypt-based stem cells localized near the base of the crypts. As developing IECs progress up the crypt-villus axis, there is a cessation of proliferation and subsequent differentiation into various cell types (e.g., enterocytes, goblet, Paneth cells) that work synergistically to mediate GI-specific functions in digestion, barrier, and immunity (Allaire *et al*, 2018).

Among those functionally distinctive epithelial cell types, tuft cells are not a newly discovered cell type, but their function was poorly understood until recently. Key roles of tuft cells include participation in chemosensing, type 2 immunity, as well as cholinergic signaling in the gut (Grencis & Worthington, 2016; Gerbe & Jay, 2016; Banerjee *et al*, 2018). Thus, tuft cells have been shown to enhance epithelial integrity and are essential for protecting the host during enteric infections and inflammatory reactions (Qu *et al*, 2015; May *et al*, 2014). Mechanisms that regulate proliferation and differentiation of IECs such as tuft cells are tightly orchestrated to ensure proper maintenance of the GI tract. Still, the molecular and cellular factors that regulate intestinal homeostasis remain largely unclear.

Inflammatory bowel disease (IBD), consisting mainly of Crohn’s disease and ulcerative colitis is a chronic inflammatory disorder of the GI tract caused by multiple factors (Baumgart & Carding, 2007). Several factors are involved in the pathogenesis of IBD, including the presence of IBD susceptibility genes, altered microbial flora, excessive innate/adaptive immunity, defective autophagy, and reduced mucosal epithelial barrier defense (Graham & Xavier, 2020; Ananthakrishnan *et al*, 2018). IBD causes dysregulation of many pathways involved in the maintenance of intestinal barrier integrity such as proliferation, migration, differentiation, and cell death. Excessive tissue injury and inadequate regeneration further lead to an increased risk of developing diseases such as cancer (Aardoom *et al*, 2018; Nadeem *et al*, 2020).

Inositol polyphosphate multikinase (IPMK) is an enzyme with broad substrate specificity that catalyzes the production of inositol polyphosphates (for example, inositol 1,3,4,5,6-pentakisphosphate) and phosphatidylinositol 3,4,5-triphosphates (Saiardi *et al*, 1999; Anutosh Chakraborty *et al*, 2011). In addition to its catalytic role critical for inositol phosphate metabolism, IPMK noncatalytically regulates major signaling factors including mechanistic target of rapamycin (mTOR), adenosine 5′-monophosphate–activated protein kinase (AMPK), p53, and serum response factor (SRF) (Lee *et al*, 2021). Collectively, IPMK acts as a signaling hub in mammalian cells that coordinates the activity of various signaling networks (Kim *et al*, 2017a). Recently, genome-wide association studies have revealed IPMK as a putative risk gene for inflammatory bowel disease (IBD) (Yokoyama *et al*, 2016; O’Donnell *et al*, 2019). Furthermore, a germ-line mutation in *Ipmk* was discovered in patients with familial small intestinal carcinoids, suggesting a role of IPMK in the homeostatic control of intestinal tissues (Sei *et al*, 2015). Accordingly, we asked whether intestinal epithelial IPMK may also play a critical role in establishing and maintaining tissue homeostasis in the gut. In this report, we characterize the effects of conditionally ablating IPMK in intestinal epithelial cells on the colonic response to dextran sulfate sodium (DSS)-induced colonic injury, a model that reproduces some features of human IBD. Our results demonstrate that IPMK-deficient colon is more vulnerable to experimental colitis, and intestinal epithelial cell-specific IPMK deficiency results in significant defects in tuft cell development. Particularly, choline acetyltransferase (ChAT) expression was significantly decreased, implying insufficient cholinergic input from tuft cells to stem cells under IPMK depletion may underlie exacerbated colitis phenotype.

## Results

### IPMK deficiency in colonic epithelium exacerbates experimental colitis and attenuates recovery

Based on the previous reports identifying several IPMK gene mutations as risk SNPs for human inflammatory bowel disease, we sought to test whether IPMK plays a significant role either in intestinal homeostasis or damaged and inflaming situation that resemble human IBD. To define the involvement of IPMK specifically in the intestinal epithelium a IPMK^ΔIEC^ (*Ipmk*^f/f^; *Villin-Cre*) mouse line was generated to achieve intestinal epithelial cell-specific IPMK depletion **(Figure S1A)**. Both IPMK transcript and protein were verified as successfully depleted in intestinal epithelia, small intestine, and colon. IPMK was unaffected in other organs **(Figure S1B, S1C)**.

Despite IPMK SNPs being repeatedly found in IBD patients, IPMK^ΔIEC^ mice did not show a significant difference in body weight compared to IPMK^f/f^ control mice in homeostatic conditions **(Figure S1D)**. Assessment of intestinal permeability using FITC-dextran also revealed that IPMK-deficient intestinal epithelium has no critical defect in its barrier function **(Figure S1E)**. In order to dissect the impact of IPMK depletion on the cell level, epithelial cells were isolated from mouse colonic tissues and analyzed. In contrary to previous findings in mouse embryonic fibroblasts (Maag *et al*, 2011), IPMK-deficient epithelial cells appeared indifferent to wild-type cells in terms of both AKT phosphorylation and mTORC1 downstream signaling cascades **(Figure S1F)**. These results collectively suggested that the deletion of IPMK in intestinal epithelium does not cause critical functional abnormalities in a normal, healthy gut.

Some factors, such as Regnase-1 (Nagahama *et al*, 2018), become prominent in intestinal homeostasis only when the tissue is damaged or inflamed. Therefore, we aimed to analyze IPMK^ΔIEC^ mice in pathological conditions. A DSS-induced acute colitis model was chosen to represent human IBD **(Figure 1A)**, and physiological features of diseased mice were assessed as in normal status. Surprisingly, IPMK^ΔIEC^ mice exhibited significantly attenuated recovery from colitis. While IPMK^f/f^ mice almost reached their initial body weight at day 17 (after 12 days post DSS severance), IPMK^ΔIEC^ mice still retained colitis-induced weight loss **(Figure 1B)**. The gut epithelial barrier was also much more permeable in IPMK^ΔIEC^ mice, as indicated by higher FITC-dextran levels in IPMK^ΔIEC^ mice sera **(Figure 1C)**. Histological analyses of colonic inflammation and damage on crypt structure further confirmed that IPMK deficiency in colonic epithelium results in exacerbated colitis upon acute DSS administration **(Figure 1D)**. This observation led us to compare the inflammation signature of wild-type and IPMK-deficient colonic epithelium. Interestingly, expression levels of inflammatory cytokines involved in colitis were not significantly different between IPMK^ΔIEC^ and IPMK^f/f^ mice **(Figure S2A)**. Inflammatory signaling pathways including NF-κB were unchanged, in parallel with cytokine levels **(Figure S2B)**.

**Figure 1.**
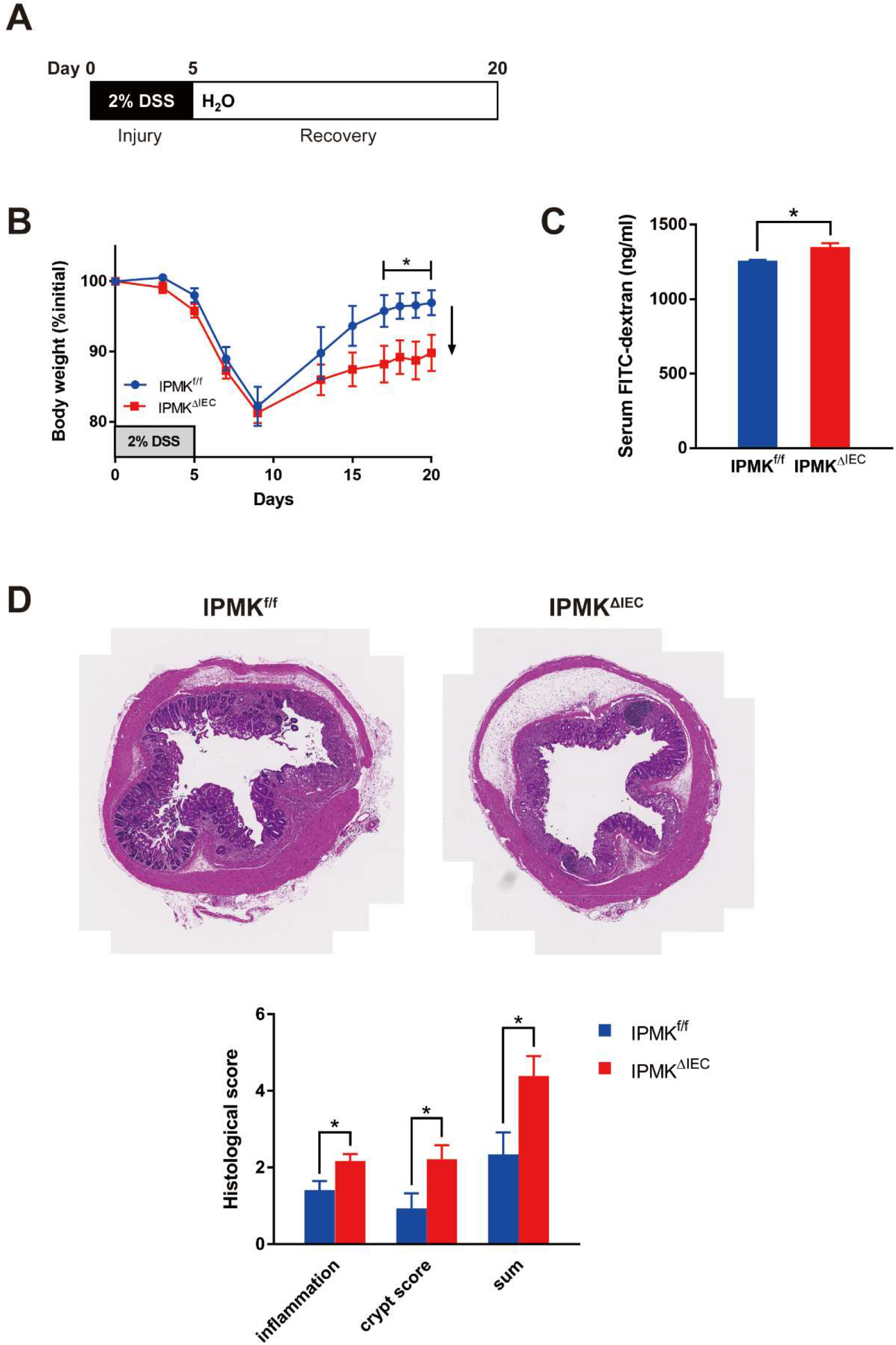
IPMK deficiency in colonic epithelium exacerbates experimental colitis and attenuates recovery. (A) Experimental scheme of DSS-induced acute colitis model. 2% DSS was given for 5 days and exchanged for normal water for recovery. Mice were sacrificed at day 8 for analysis of actively inflaming tissue or at day 20 for end-point analysis. (B) IPMK^ΔIEC^ mice exhibited attenuated recovery from colitis. Body weight was monitored at indicated time points and calculated as the percentage of initial body weight (*n*=8, 7 per genotype). (C) FITC-dextran permeability of the colonic wall was higher in IPMK^ΔIEC^ mice at day 20. The concentration of FITC-dextran in mouse serum was measured 4 hr after 200 mg/kg given by oral gavage (*n*=8, 6 per genotype). (D) IPMK-deficient colons showed exacerbated tissue damage at day 8. Representative H&E staining images of distal colon cross-sections. Histological scoring was done in two categories (inflammation and crypt score) and combined (*n*=8, 11 per genotype). \**P* < 0.05.

### IPMK-deficient colon shows significantly decreased tuft cell number

Since we could not find marked differences in inflammation in IPMK-deficient colons, transcriptomic analysis was performed to elucidate what underlies the severe disease in IPMK^ΔIEC^ mice. Unexpectedly, the top-ranked category of differentially expressed genes (DEG) was ‘keratinization’, a seemingly colitis-irrelevant biological process **(Figure 2A)**. Those genes included members of the small proline-rich (Sprr) protein family, such as *Sprr2d* and *Sprr2h*, and appeared to be dramatically increased in the IPMK-deficient colon **(Figure 2B)**. Normally, those genes act as key components of keratinocyte differentiation and are strongly induced upon differentiation of stratified squamous epithelia (epidermis) (Tesfaigzi & Carlson, 1999). Significant upregulation of the *Sprr* gene family in the colon can be interpreted in two ways. First, *Sprr2h* mRNA level was reported to be increased in an experimental colitis model, possibly as a compensatory action for deteriorated barrier function (Fang *et al*, 2012), so higher *Sprr* levels may reflect functionally impaired, ‘leaky’ colonic epithelia. Secondly, keratinocyte differentiation marker genes such as *Sprr2a* and *Krt10* are known to be regulated by Pou2f3 (Ma *et al*, 2020), a transcription factor that is abundantly expressed in the epidermis. However, in the colon, Pou2f3 is a cell fate-determining factor that is indispensable for specification of tuft cells (Yamashita *et al*, 2017), a rare and functionally distinctive population of the intestinal epithelium. We speculated that differences in Pou2f3 expression could underlie this keratinization-like transcriptomic profile, and surprisingly, Pou2f3 was significantly downregulated in IPMK^ΔIEC^ mice along with other tuft cell markers such as *Trpm5* and *Gnat3* **(Figure 2C, 2D)**. Other epithelial cell populations that are represented by *Anpep* (colonocytes), *Muc2* (goblet cells), *Chga* (enteroendocrine cells), and *Reg4* (deep crypt secretory cells), did not exhibit significant variation. So, this concurrent reduction in tuft cell-associated genes strongly implicates the alteration of tuft cell populations in the IPMK-deficient colon.

**Figure 2.**
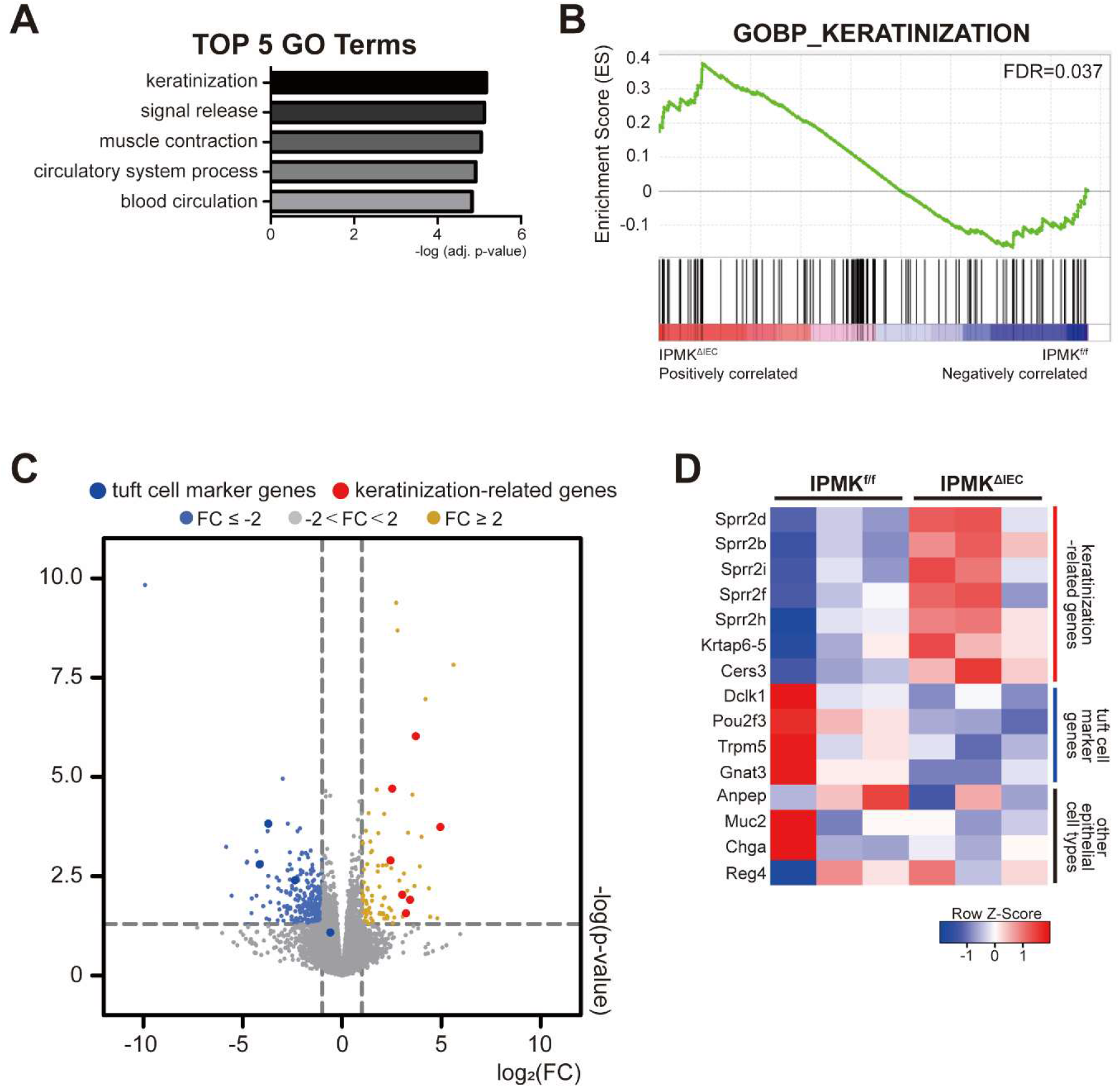
Transcriptomic profile of IPMK-deficient colon reveals gene signatures associated with keratinization and tuft cell alteration. (A) The top 5 GO terms were calculated using adjusted p-values. (B) Enrichment set data showed significant changes of keratinization terms by GSEA. (C) Volcano plot of the DEG marked with tuft cell markers and keratinization-related genes. (D) Heatmap representing selected DEGs. Genes associated with keratinization were upregulated whereas tuft cell markers were downregulated, and other epithelial cell type-related genes did not show significant alterations. Mice were sacrificed on day 20 for RNA sequencing of colonic tissues.

Accordingly, decreased mRNA expression of tuft cell markers was further validated through qRT-PCR in colonic epithelial cells. Earlier studies reported *Dclk1, Pou2f3, Trpm5*, and *Gnat3* as tuft cell-specific genes in intestinal epithelia, and expression levels of all three genes except for Trpm5 showed significant decline in IPMK^ΔIEC^ mice **(Figure 3A)**. In addition, Dclk1, a marker protein of colonic tuft cells, was upregulated at the protein level. These results clearly emphasized the need for precise quantification of tuft cell populations either in wild-type and IPMK-deficient colon under colitis **(Figure 3B)**. Therefore, we directly analyzed the relative abundance of tuft cells using flow cytometry, assessing Siglec F- and EpCAM-double positive cells in isolated colonic epithelia. The percentage of tuft cells in total colonic epithelial cells was significantly lower in IPMK^ΔIEC^ mice, only about 70% of the control group **(Figure 3C)**. This suggests the alteration in colonic tuft cell populations is the main cause of exacerbated colitis in IPMK^ΔIEC^ mice upon acute DSS administration.

**Figure 3.**
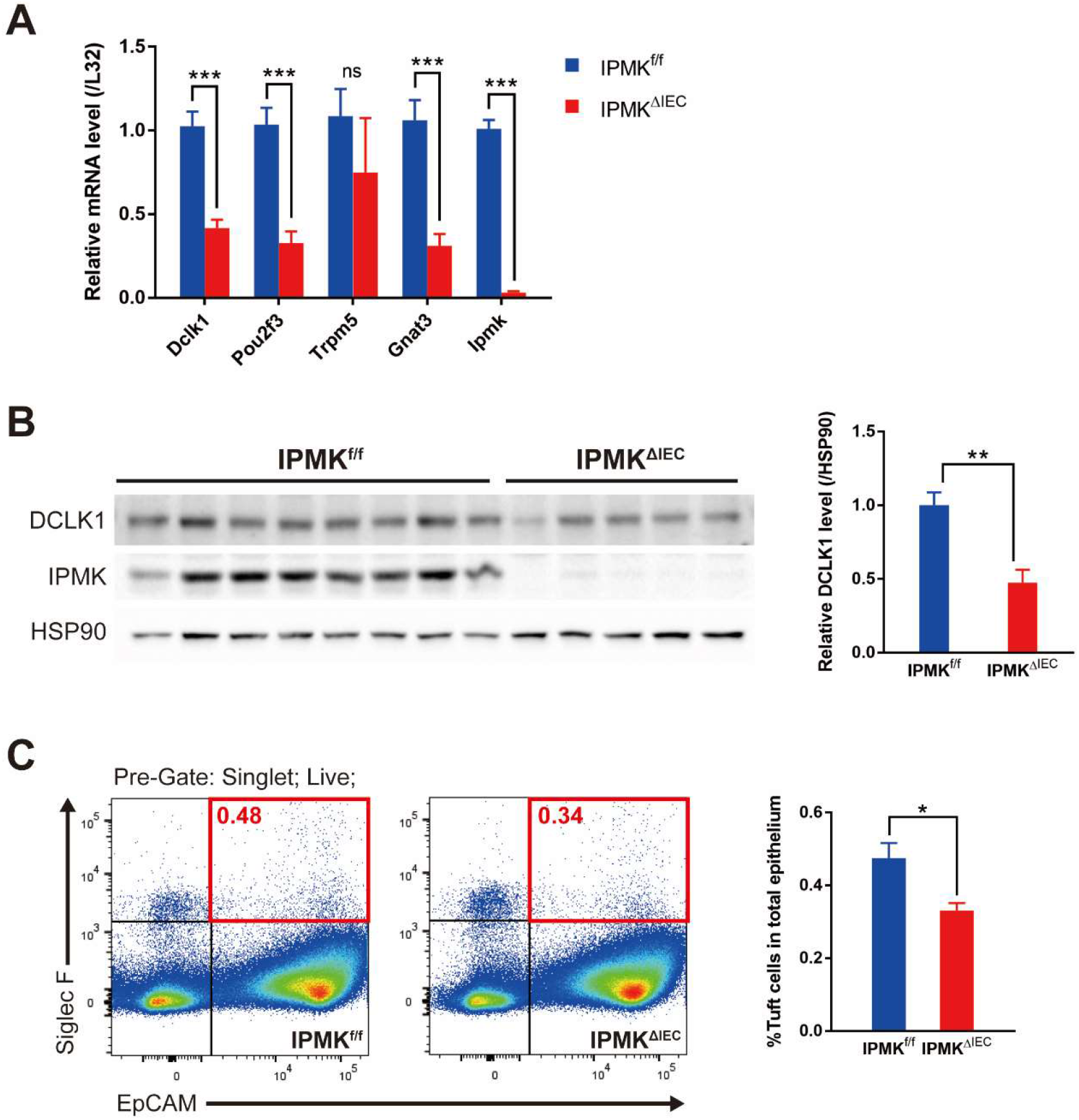
Colonic IPMK depletion leads to significantly decreased tuft cell numbers upon acute colitis. (A) Colonic epithelial cells from IPMK^ΔIEC^ mice had decreased expression of tuft cell marker genes. Each mRNA was assessed by qRT-PCR and normalized to the expression of the L32 gene (*n*= 8, 5 per genotype). (B) DCLK1 protein level was lower in IPMK-deficient colon as shown in representative blots. Band intensities were quantified by densitometry using ImageJ (*n*= 8, 5 per genotype). (C) Tuft cell abundance in the colonic epithelium was quantified by flow cytometry. Epithelial cells were defined by EpCAM expression, and Siglec F-positive epithelial cells (red square) were considered as tuft cells (*n*= 8, 5 per genotype). All mice were sacrificed at day 8.\**P* < 0.05, \*\**P*<0.01.

### IPMK regulates cholinergic input to stem cells via governing tuft cell development

To investigate the relationship between IPMK deletion and tuft cell decrease, we first tested whether the occurrence of this phenomenon is limited to the pathologic condition. In normal unchallenged adult mice of 10 weeks-old, the abundance of each colonic epithelial cell type was analyzed using marker genes. *Dclk1* of tuft cells was the only gene that was significantly decreased in IPMK^ΔIEC^ mice, while others did now show a significant difference between groups **(Figure 4A)**. Therefore, only tuft cells are specifically affected by the depletion of IPMK in the intestinal epithelium. The protein level of Dclk1 was also reduced in the IPMK-deficient colon at homeostasis **(Figure 4B)**, and direct measurement of Siglec F- and EpCAM-double-positive cells using flow cytometry again validated the reduction of tuft cell numbers in healthy IPMK^ΔIEC^ mice **(Figure 4C)**. Collectively, these findings suggest that depletion of IPMK in colonic epithelium already impaired tuft cell development before colitis progression, and these defective tuft cells subsequently result in the vulnerability of IPMK^ΔIEC^ mice upon DSS administration, without any notable abnormalities in homeostatic condition.

**Figure 4.**
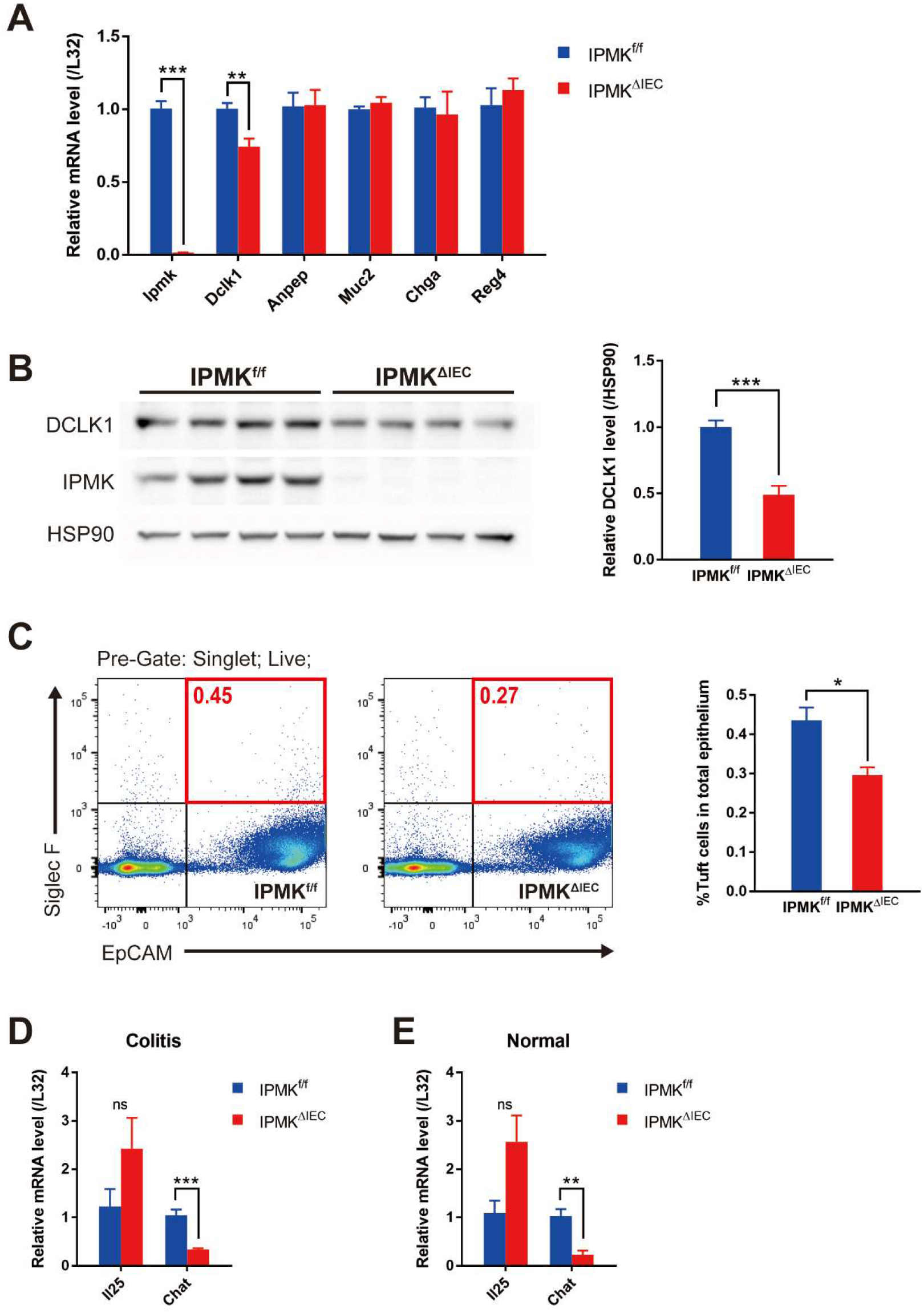
IPMK governs tuft cell development and regulates acetylcholine production in the colonic epithelium. (A) Basal expression of colonic epithelial cell lineage-specific marker genes was analyzed by qRT-PCR and normalized to the expression of the L32 gene. Only Dclk1, a tuft cell marker, appeared to be downregulated (*n*=6, 3 per genotype). (B) Normal colonic epithelial cells showed decreased DCLK1 protein. Band intensities of representative blots were quantified by densitometry using ImageJ (*n*= 4, 4 per genotype). (C) The percentage of tuft cells in the normal healthy colonic epithelium was quantified by flow cytometry and was lower in IPMK^ΔIEC^ mice. EpCAM- and Siglec F-double-positive cells (red square) were counted as tuft cells (*n*= 4, 4 per genotype). (D) On day 8, ChAT expression was reduced in IPMK-deficient colonic epithelial cells upon colitis while IL-25 expression level was not altered. qRT-PCR results were normalized to the L32 gene (*n*= 7, 5 per genotype). (E) ChAT mRNA was also downregulated in IPMK^ΔIEC^ even in normal conditions (n= 4, 4 per genotype). \**P* < 0.05, \*\**P*<0.01, \*\*\**P*<0.001

**Figure 5.**
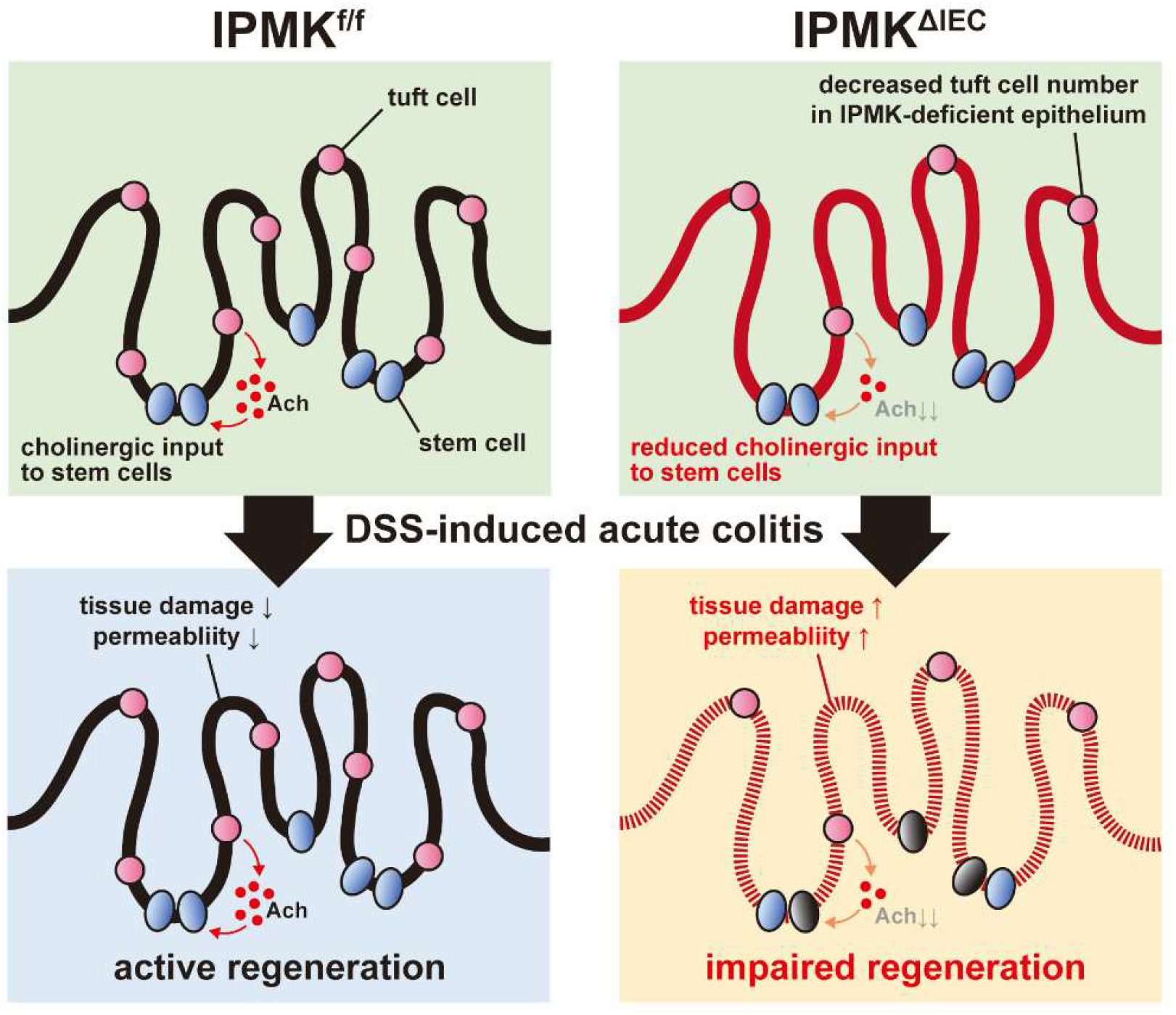
A proposed model of IPMK actions in the gut. The number of tuft cells, providing acetylcholine that supports intestinal stem cell function, is reduced in the IPMK-deficient colon. Insufficiency in tuft cells is tolerated in the normal homeostatic epithelium, so IPMK^ΔIEC^ mice do not show any apparent abnormalities. However, when acute damage is applied to a colon with DSS-induced experimental colitis, IPMK-deficient colons exhibits impaired regeneration, which may be due to decreased cholinergic input to stem cells, thereby exacerbating the colitis phenotype.

Currently, tuft cells are known to be involved in intestinal biology by two distinctive ways; as a regulator of type 2 immune response (Howitt *et al*, 2016; Gerbe *et al*, 2016) and a paracrine supporter of intestinal stem cells (Middelhoff *et al*, 2017). Those functions are mostly mediated by IL-25 and acetylcholine respectively, that are secreted by tuft cells. To elucidate how reduced tuft cells in IPMK-deficient colon worsen the DSS-induced acute colitis, the mRNA expression level of IL-25 and ChAT (choline acetyltransferase; a key enzyme for acetylcholine synthesis in colonic tuft cells) was compared between IPMK^f/f^ and IPMK^ΔIEC^ mice. Both in normal and pathologic states, ChAT expression appeared significantly lower in IPMK-deficient colon **(Figure 4D, 4E)**, indicating cholinergic input for intestinal stem cells is decreased.

## Discussion

To summarize, intestinal epithelia-specific IPMK deletion drives the mouse colon to be more prone to DSS-induced acute colitis. Unlike our expectations from previous studies on IPMK actions in PI3K and inflammation signaling, IPMK deletion made no impact on intestinal epithelial growth signaling (e.g. PI3K-AKT-mTOR) or NF-κB inflammatory activation. Instead, to our surprise, Tuft cells, which normally promote intestinal stem cell proliferation by secreting acetylcholine, were significantly reduced in the IPMK-deficient colon. As a result, recovery of damaged tissue was attenuated, likely due to an insufficient amount of acetylcholine. This means that IPMK normally protects intestinal epithelium from damage induced by acute inflammation, and through a yet undefined mechanism, IPMK governs the specification of undifferentiated epithelial cells into tuft cells.

Previous studies demonstrated that tuft cell ablation is tolerated under normal homeostatic conditions, but exhibits impaired regeneration of colonic tissues upon acute damage such as irradiation (May *et al*, 2014) and DSS-induced acute colitis (Qu *et al*, 2015) in accordance with our study. Defective tissue repair derived from tuft cell insufficiency was fully restored by administration of M3 muscarinic receptor agonist, replacing endogenous cholinergic input for intestinal stem cells (Hayakawa *et al*, 2017). Therefore, to prove that diminished acetylcholine release from tuft cells is responsible for IPMK^ΔIEC^ mice phenotype, a similar approach using a specific acetylcholine receptor agonist should be employed in future studies.

Until now, two factors are known to be indispensable for the development of tuft cells; *Atoh1* and *Pou2f3*. Whereas *Atoh1* is commonly required for differentiation of all secretory lineage, *Pou2f3* is specific to tuft cells (Gerbe *et al*, 2011). The fact that tuft cells were the only colonic epithelial cell population impacted by the absence of IPMK suggests that IPMK does not influence the *Atoh1*-dependent specification processes. Therefore, IPMK is likely to act downstream of *Atoh1* but upstream of *Pou2f3*. Considering that IPMK serves not only as a kinase that produces PIP_3_, IP_4_, and IP_5_ but also as a regulator of diverse nuclear events in an enzymatically independent fashion, how IPMK controls tuft cell differentiation should be further investigated.

Tuft cells have been actively studied in the small intestine, rather than the colon, focusing specifically on its role as type 2 immune response regulator. Interestingly, small intestinal tuft cells are clearly distinguished from colonic tuft cells because succinate, a metabolite produced by helminths, triggers dramatic expansion of small intestinal tuft cells while colonic tuft cells remain unchanged (Nadjsombati *et al*, 2018; Lei *et al*, 2018). This demonstrates that the specification mechanism of tuft cells can vary between different organs. Therefore, determining the involvement of IPMK in tuft cell development in other organs possessing tuft cells (e.g. trachea, lung, thymus, etc) would be beneficial. A recent single-cell RNA sequencing study of small intestinal epithelium discovered that tuft cells are transcriptomically divided into two groups; one population that primarily expresses inflammation-related genes and the other expresses neuromodulatory genes (Haber *et al*, 2017). In contrast to the small intestine, colonic tuft cells have not yet been fully analyzed at the single-cell level. In this study, we observed that IPMK specifically reduces ChAT expression without affecting the IL-25 level. Whether this result indicates that colonic tuft cells are also distinguished into two functionally distinct groups and IPMK depletion specifically affects only one group should be further analyzed.

Several clinical studies have independently reported that reduced tuft cell numbers in the intestinal tract are associated with ileal Crohn’s disease (Banerjee *et al*, 2020) and quiescent ulcerative colitis (Kjærgaard *et al*, 2021), suggesting that insufficient tuft cells are closely related to the pathology of inflammatory bowel diseases. Since we newly propose a IPMK-tuft cell-IBD axis in this study, targeting IPMK has the potential to be a novel therapeutic strategy to treat IBD.

## Materials and methods

### Mice

Animal protocols were performed in accordance with the guidelines approved by the Korea Advanced Institute of Science and Technology Animal Care and Use Committee. Intestinal epithelium-specific IPMK knockout mice were generated by crossing *Ipmk-* floxed (Kim *et al*, 2017b) and *Villin-Cre* mice (#004586 B6.Cg-Tg(Vil1-cre)997Gum/J, The Jackson Laboratory). Mice were backcrossed to C57BL/6J for at least 5 generations. 8 to 20 weeks-old mice (both male and female) were used in this study. A single experimental cohort consisted of age and sex-matched mice, of which age does not differ by more than 2 weeks. Mice used in this study were maintained in a specific pathogen-free facility at KAIST Laboratory Animal Resource Center.

### DSS-induced acute colitis model

Dextran sulfate sodium salt (colitis grade, M.W. 36,000-50,000 Da, MP Biomedicals) was given to mice as a 2% filtered solution in drinking water, ad libitum, to induce acute colitis. DSS-containing water was refreshed once every 2 to 3 days. After 5 days of DSS administration, mice were allowed to recover with normal drinking water. Body weight and survival of each mouse was monitored at the indicated time points. No more than 5 mice were kept in one cage.

### FITC-dextran intestinal permeability assay

Mice were fasted (water and food) for 4 hours, and then orally gavaged with Fluorescein isothiocyanate (FITC)-dextran (FD4, Sigma-Aldrich) solution in DPBS using a 400 mg/kg dose. For colitis-induced mice, a 200 mg/kg dose was used. After 4 hours, mice were provided with water *ad libitum*, and 100 μl of blood was drawn from the tail. Serum was prepared by incubating at room temperature for 15 minutes then centrifuging at 4°C, 1,500×g for 10 minutes. Serum samples (diluted in DPBS) from each mouse were loaded as technical replicates on black plates along with known concentrations of FITC-dextran standards, and fluorescence intensity was measured (excitation 485nm, emission 535nm) using a Multidetection Microplate Reader (TriStar^2^ LB 942, BERTHOLD Technologies).

### Histology

Colonic tissues were harvested from mice after euthanasia. Fecal contents were removed by repeated flushing with pre-chilled PBS. A distal region (∼1 cm) was cut and placed in an Ep-tube to make cross-sectional histology samples. Otherwise, colons were opened longitudinally along their mesenteric line and made into swiss-rolls. In both cases, tissues were then fixed in 10% neutral buffered formalin overnight at room temperature. Subsequent processing and H&E staining were conducted by KPNT (Cheongju, Korea), an inspection institution. Bright-field images of stained slides were obtained by Axio Scan Z1 Slide Scanner (Carl Zeiss).

### Intestinal epithelial cell isolation

Colonic epithelia were isolated according to the protocol of Ma group (Shao *et al*, 2013) with some modifications. Briefly, excised and feces-cleared mouse colons were opened along the mesenteric line, cut again into longitudinal halves, and cut into 2-3 mm sized pieces. Tissue fragments were gently agitated in ice-cold HBSS supplemented with 2% glucose and 1 mM DTT at 4°C for 10 minutes in order to eliminate mucus. Tissues were then moved into DPBS (Ca^2+^ and Mg^2+^-free) containing 10 mM EDTA and incubated for 15 minutes at 37°C with 200 rpm shaking. After robust vortexing, colonic epithelia were released into the supernatant. Epithelium-containing supernatants were passed through a 100 µm cell strainer, and cells were collected by centrifugation at 4°C (1,000×g, 5 minutes). For flow cytometry, the epithelium releasing step was repeated 3 more times without further incubation at 37°C. Supernatants were combined and passed through a 70 µm cell strainer, collected by centrifugation, and washed with ice-cold HBSS. After centrifugation, cell pellets were resuspended in pre-warmed HBSS containing 1 U/ml Dispase (07913, Stemcell Technologies) and 10 µg/ml DNase I (11284932001, Roche), and incubated for 10 minutes at 37°C with 200 rpm shaking to yield a single-cell suspension. Vortexed again, cells were passed through a 40 µm cell strainer and collected by centrifugation at 4°C (500×g, 5 minutes). Cell pellets were washed with ice-cold DPBS once and counted.

### Tuft cell quantification using flow cytometry

Colonic epithelial cells were stained with eBioscience™ Fixable Viability Dye eFluor™ 506 (65-0866-14, Invitrogen) in a 1:1000 ratio for dead cell exclusion. Cells were Fc-blocked using Purified Rat Anti-Mouse CD16/CD32 (Clone 2.4G2, 553142, BD Pharmingen™) for 10 minutes and washed with FACS buffer (Ca^2+^ and Mg^2+^-free DPBS supplemented with 0.5% BSA and 0.3% sodium azide). For surface marker staining, PE Rat Anti-Mouse Siglec-F (Clone E50-2440, 552126, BD Pharmingen™) antibody and APC anti-mouse CD326 (EpCAM) antibody (clone G8.8, 118214, Biolegend) were used in a 1:50 and 1:100 dilution ratio, respectively. Antibody staining was performed for 30 minutes. All procedures were conducted at 4°C in the dark. Tuft cells were acquired by BD LSRFortessa (BD Biosciences) and analyzed using FlowJo v10.7.2 (Tree Star). The percentage of tuft cells was calculated by dividing the number of EpCAM- and PE- double-positive cells by the number of EpCAM-positive cells.

### Immunoblotting

After flushing, colons were cut into appropriate sizes and mechanically homogenized in RIPA buffer (50 mM Tris-HCl pH 7.4, 150 mM NaCl, 1 mM EDTA, 1% NP-40, 0.25% sodium deoxycholate, 0.1% sodium dodecyl sulfate, 2 mM PMSF, 10 mM sodium fluoride, 2 mM sodium orthovanadate, 1 mM sodium pyrophosphate, 1X protease inhibitor cocktail) using TissueRuptor (Qiagen). Isolated epithelial cells were also lysed in RIPA buffer. Following incubation for 30 minutes at 4°C, lysates were clarified by centrifugation at 13,000 rpm for 20 minutes, and protein concentration was determined using Pierce™ BCA Protein Assay Kit (23225, Thermo Scientific™) according to the manufacturer’s protocol. For immunoblotting, antibodies against the following proteins were purchased from the indicated sources: DCLK1 (ab31704) (Abcam); HSP90 (sc-13119) (Santa Cruz Biotechnology); α-tubulin (T5168) (Sigma-Aldrich); phospho-S6 (5364), S6 (2217), phospho-AKT T308 (4056), phospho-AKT S473 (9271), AKT (9272), phospho-NFκB (3033), NFκB (8242), phospho-JNK (4668), β-actin (4970) (Cell Signaling Technology). Anti-mouse IPMK antibody was a custom rabbit polyclonal antibody from Covance raised against a synthetic peptide starting with Cys followed by mouse IPMK amino acids 295 to 311 (SKAYSRHRKLYAKKHQS) (Maag *et al*, 2011). Densitometry analysis of blots was performed in ImageJ 1.53e.

### RNA extraction and qRT-PCR

Total RNA was extracted from mouse tissues and epithelial cells using TRI-Reagent (TR-118, Molecular Research Center) or RNeasy Lipid Tissue mini kit (74804, Qiagen) according to the manufacturer’s protocol. A maximum of 3 µg of total RNA was reverse-transcribed to yield first-strand complementary DNA with SuperiorScript III Reverse Transcriptase (RT006L, Enzynomics). Quantitative PCR analyses were performed on a StepOnePlus Real-Time PCR System (Applied Biosystems) using SYBR Green (QPK-201, Toyobo) and 20 or 25 ng of cDNA. The relative expression level of each target gene was normalized to *L32* gene using ΔΔC_t_ method. Primer sequences for qRT-PCR are provided in the table below (all *Mus musculus*).

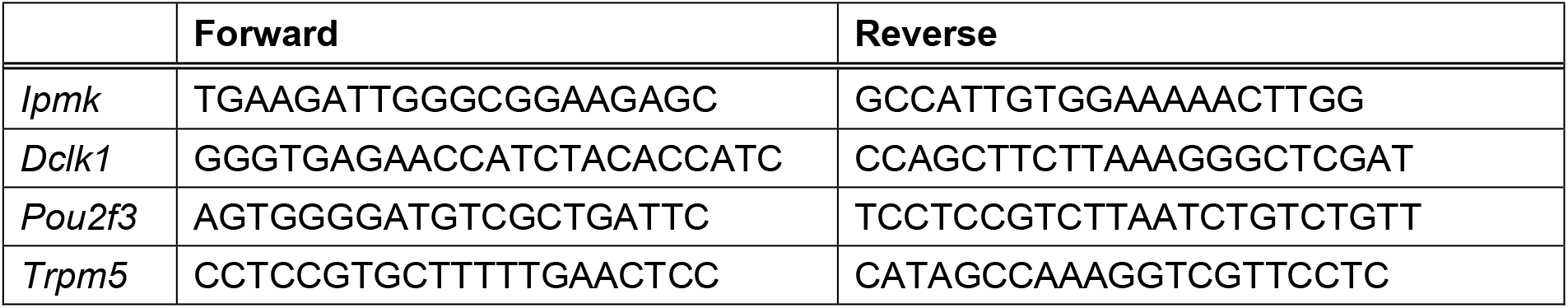
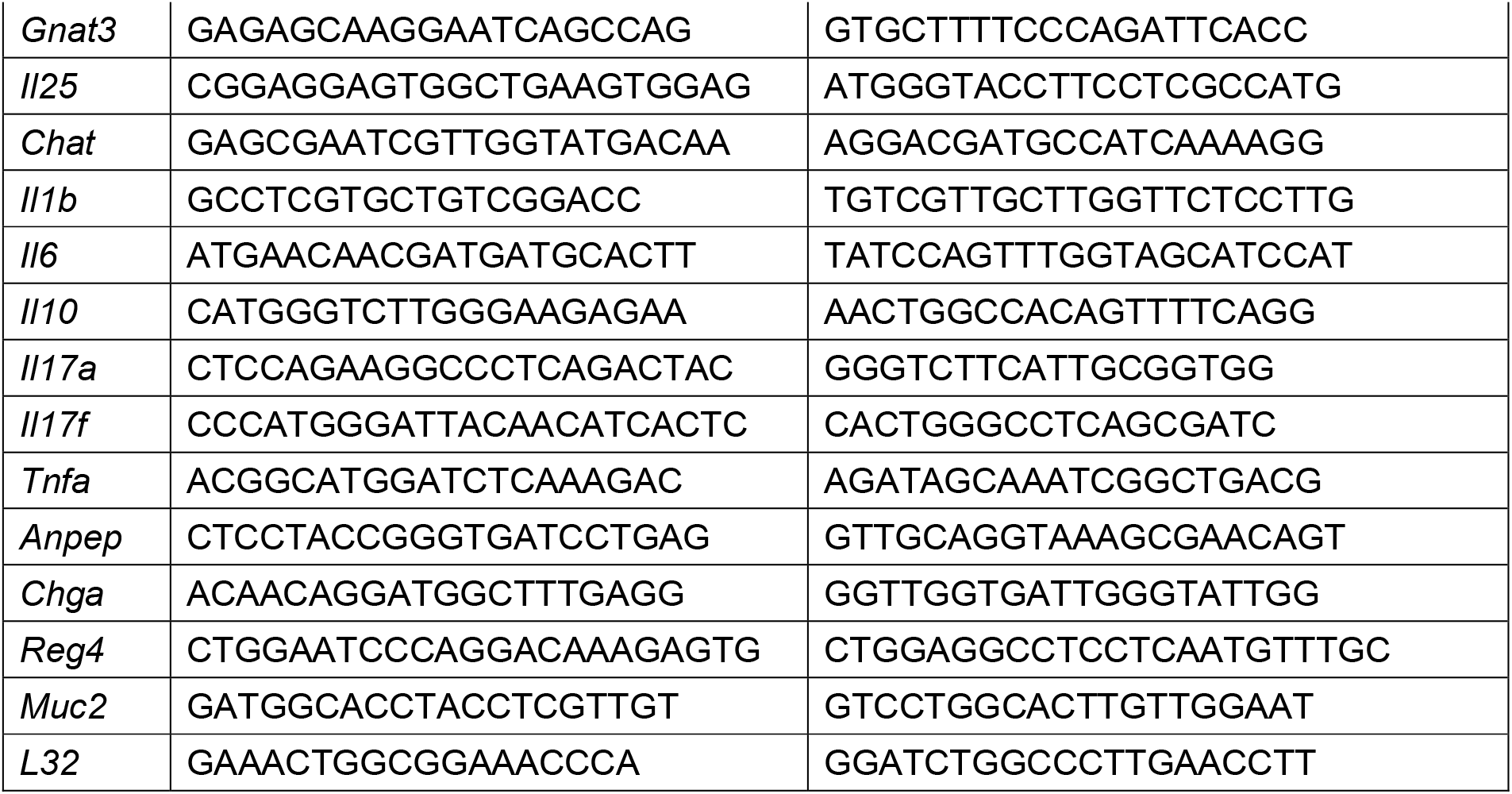

### mRNA sequencing and data processing

Library preparation, sequencing, preprocessing, genome mapping, and gene expression quantification were conducted by LAS Inc. (http://las.kr) as demonstrated below. Total RNA was isolated from tissue using TRI-reagent-based method. Total RNA (500 ng) was processed for preparing a whole transcriptome sequencing library. Enrichment of whole transcriptome RNA by depleting ribosomal RNA (rRNA) was conducted to prepare the whole transcriptome sequencing library using MGIEasy RNA Directional Library Prep Kit (MGI) according to the manufacturer’s instruction. After the rRNA was depleted, the remaining RNA is fragmented into small pieces using divalent cations under elevated temperature. The cleaved RNA fragments are copied into first strand cDNA using reverse transcriptase and random primers. Strand specificity is achieved in the RT directional buffer, followed by second strand cDNA synthesis. These cDNA fragments have the addition of a single ‘A’ base and subsequent ligation of the adapter. The products are then purified and enriched with PCR to create the final cDNA library. The double stranded library is quantified using QuantiFluor ONE dsDNA System (Promega) and equal 330 ng in a total volume of 60 ml or less. The library was cyclized at 37 °C for 60 min, and then digested at 37 °C for 30 min, followed by cleanup of the circularization product. To make the DNA nanoball (DNB), the library was incubated at 30 °C for 25 min using DNB enzyme. Finally, the library was quantified by QuantiFluor ssDNA System (Promega). Sequencing of the prepared DNB was conducted using the MGIseq system (MGI) with 100 bp paired-end reads.

Potentially existing sequencing adapters and raw quality bases in the raw reads were trimmed by Skewer ver 0.2.2 (Jiang *et al*, 2014). The cleaned high-quality reads were mapped to the reference genome by STAR ver 2.5 software (Dobin *et al*, 2013). Since the sequencing libraries were prepared strand-specifically by using Illumina’s strand-specific library preparation kit, the strand-specific library option, --library-type=fr-firststrand was applied in the mapping process.

To quantify the mapped reads RSEM ver 1.3.0 (Li & Dewey, 2011) and default options were used. The gene annotation of the reference genome mm10 from GENCODE genome (https://www.gencodegenes.org) in GTF format was used as gene models. The differentially expressed genes between the two selected biological conditions were analyzed by DESeq2 ver 1.26.0 software (Love *et al*, 2014). To compare the expression profiles among the samples, the normalized expression values of the selected a few hundred of the differentially expressed genes were clustered (unsupervised) by in-house R scripts. The scatter plots for the gene expression values and the volcano plots for the expression-fold changes and p-values between the two selected samples were drawn by in-house R scripts.

### Analysis and visualization of mRNA-sequencing data

Gene ontology enrichment analysis (GO; http://geneontology.org/) was performed by g:Profiler ver 0.2.0 and top 5 GO biological process (GOBP) terms were visualized by −log_10_ transformed adjusted p-value. Gene set enrichment analysis (GSEA; https://www.gsea-msigdb.org/gsea/index.jsp) was performed by GSEA ver 4.1.0 with normalized count and visualized with enrichment score and calculated FDR. Volcano plots were visualized by log_2_ transformed FC and −log_10_ transformed p-value. Then tuft cell marker genes and keratinization-related genes were represented by specific plots. Heatmaps representing each term were visualized with row z-score calculated by log_2_ transformed normalized count.

### Statistical analysis

All statistical analyses were performed using Prism 7 (Graphpad Software) and all data were presented as ±SEM. Differences between groups were considered to be significant when *P*<0.05 (Student’s unpaired *t* test, two-tailed).

## Acknowledgements

We thank the members of the Fang and Kim laboratory for their assistance and advice. This work was supported by the grant from TJ Park Science Fellowship of the POSCO TJ Park Foundation (to S.E.P.) and National Research Foundation of Korea (NRF-2021R1A2C2009749 to S.F.; 2018R1A5A1024261, 2020R1A2C3005765to S.K.).

## Author Contributions

S.E.P., S.F., and S.K. conceived the work. S.E.P., S.J.P., J.R., and S.K.O. performed the experiments. S.E.P., S.J.P., S.F., and S.K. designed the experiments. S.E.P., J.J., S.J.P., J.R., S.K.O., S.L., S.F., and S.K. analyzed the data. S.E.P. and S.K. wrote the manuscript.

S.F. and S.K. supervised the study.

---

**Figure S1.**
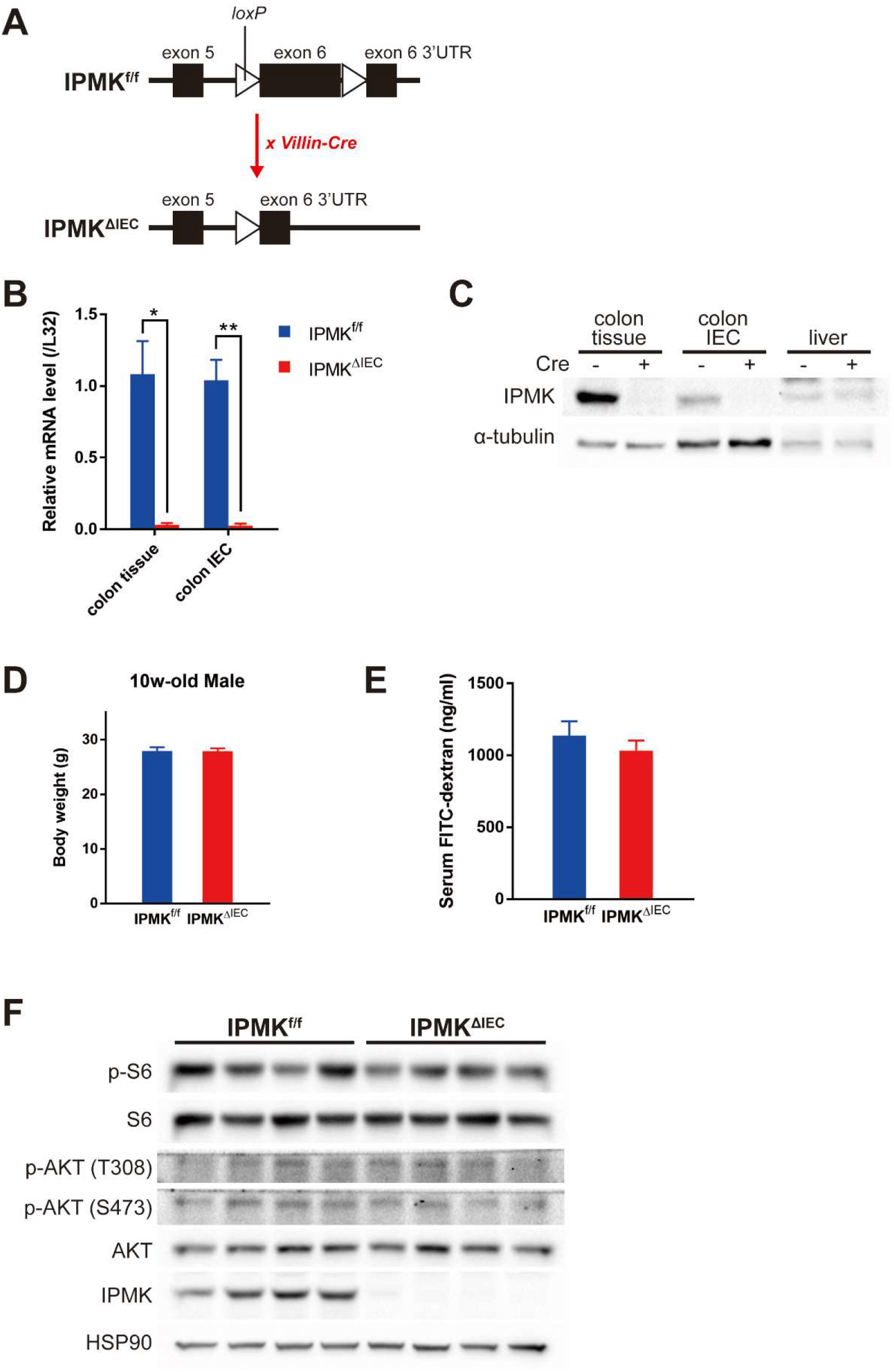
IPMK^ΔIEC^ mice appear normal in homeostatic conditions. (A) IPMK was specifically depleted in intestinal epithelium using *Villin-Cre* mice. *loxP* sequences were inserted in the exon 6 region of *Ipmk* gene, and Cre-driven deletion of flanked sequences results in truncated mRNA transcripts that are readily degraded. (B, C) Tissue-specific IPMK knockout was confirmed at the mRNA level (B) and protein level (C) (*n*=5, 3 per genotype in B). (D) Body weights of 10 week-old mice did not show any difference between genotypes (*n*=6, 3 per genotype). (E) Gut permeability did not significantly differ between genotype, in a normal state. Serum FITC-dextran concentration was measured 4 hr after 400 mg/kg dosage by oral gavage (*n*=6, 6 per genotype). (F) No notable difference was observed in blots of mTORC and AKT signaling cascade. As these blots were from a single immunoblot cohort that was used in Figure 4B, IPMK and HSP90 blots were shown again here. \**P* < 0.05, \*\**P*<0.01.

**Figure S2.**
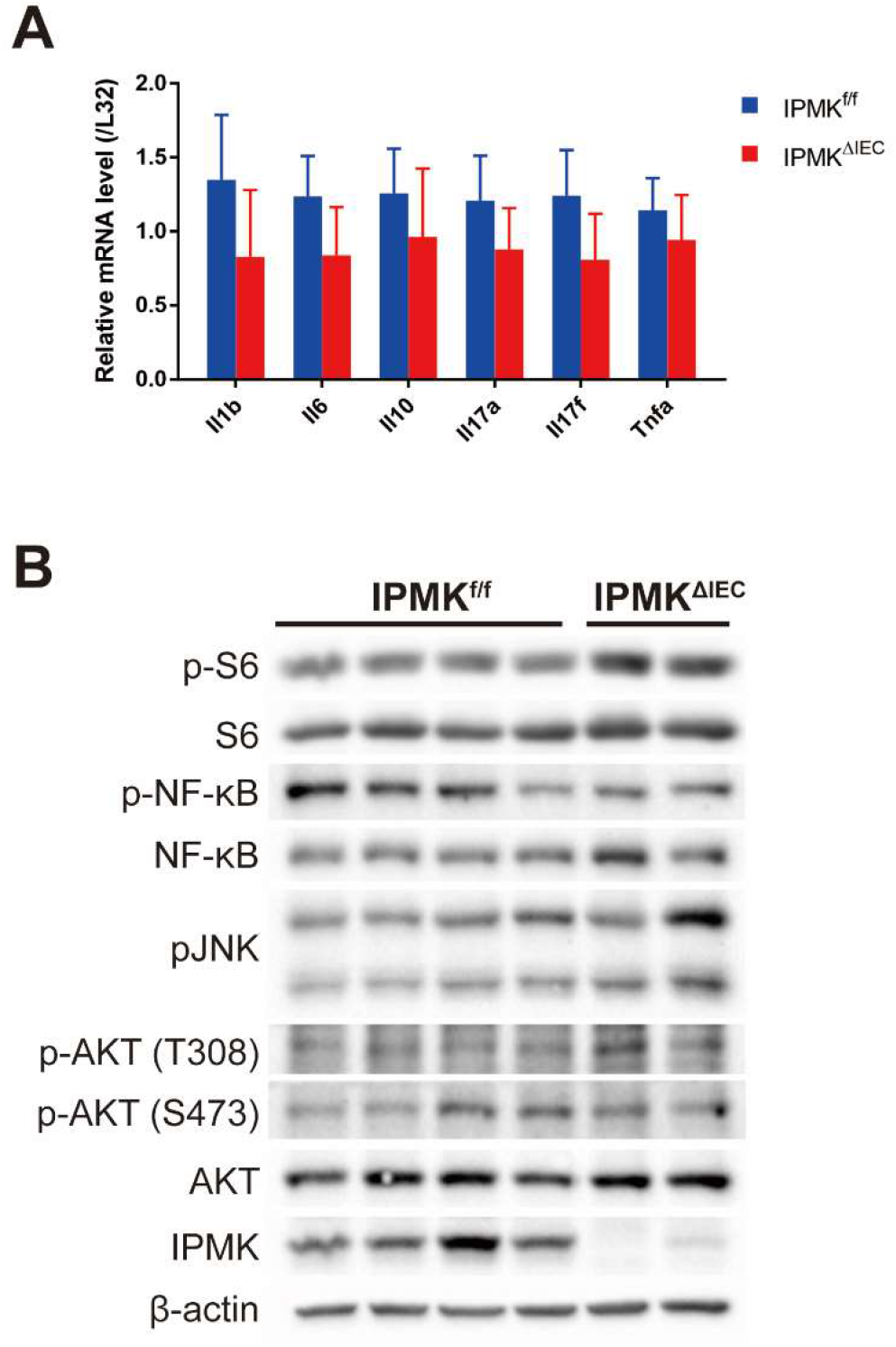
No significant difference in inflammatory signaling exists between IPMK^ΔIEC^ and control mice. (A) Although IPMK deletion exacerbates colitis phenotypes, expression of major inflammatory cytokines in colonic epithelial cells did not show any difference between genotypes (*n*=8, 5 per genotype). (B) The activity of the NF-κB pathway, regulating inflammation, and AKT signaling, governing cell survival, were assessed by immunoblotting. The phosphorylation level of key proteins was not different between IPMK^ΔIEC^ and control mice.

## References

Aardoom MA, Joosse ME, De Vries ACH, Levine A & De Ridder L (2018) Malignancy and Mortality in Pediatric-onset Inflammatory Bowel Disease: A Systematic Review. Inflamm Bowel Dis 24: 732–741

Allaire JM, Crowley SM, Law HT, Chang SY, Ko HJ & Vallance BA (2018) The Intestinal Epithelium: Central Coordinator of Mucosal Immunity. Trends Immunol 39: 677–696

Ananthakrishnan AN, Bernstein CN, Iliopoulos D, Macpherson A, Neurath MF, Ali RAR, Vavricka SR & Fiocchi C (2018) Environmental triggers in IBD: A review of progress and evidence. Nat Rev Gastroenterol Hepatol 15: 39–49

Anutosh Chakraborty, Kim S & Snyder SH (2011) Inositol Pyrophosphates as Mammalian Cell Signals. Sci Signal 4: re1

Banerjee A, Herring CA, Chen B, Kim H, Simmons AJ, Southard-Smith AN, Allaman MM, White JR, Macedonia MC, Mckinley ET, et al (2020) Succinate Produced by Intestinal Microbes Promotes Specification of Tuft Cells to Suppress Ileal Inflammation. Gastroenterology 159: 2101-2115.e5

Banerjee A, McKinley ET, Von Moltke J, Coffey RJ & Lau KS (2018) Interpreting heterogeneity in intestinal tuft cell structure and function. J Clin Invest 128: 1711–1719

Baumgart DC & Carding SR (2007) Inflammatory bowel disease : cause and immunobiology. Lancet 369: 1627–1640

Dobin A, Davis CA, Schlesinger F, Drenkow J, Zaleski C, Jha S, Batut P, Chaisson M & Gingeras TR (2013) STAR: Ultrafast universal RNA-seq aligner. Bioinformatics 29: 15–21

Fang K, Zhang S, Glawe J, Grisham MB & Kevil CG (2012) Temporal genome expression profile analysis during t-cell-mediated colitis: Identification of novel targets and pathways. Inflamm Bowel Dis 18: 1411–1423

Gerbe F, Van Es JH, Makrini L, Brulin B, Mellitzer G, Robine S, Romagnolo B, Shroyer NF, Bourgaux JF, Pignodel C, et al (2011) Distinct ATOH1 and Neurog3 requirements define tuft cells as a new secretory cell type in the intestinal epithelium. J Cell Biol 192: 767–780

Gerbe F & Jay P (2016) Intestinal tuft cells: Epithelial sentinels linking luminal cues to the immune system. Mucosal Immunol 9: 1353–1359

Gerbe F, Sidot E, Smyth DJ, Ohmoto M, Matsumoto I, Dardalhon V, Cesses P, Garnier L, Pouzolles M, Brulin B, et al (2016) Intestinal epithelial tuft cells initiate type 2 mucosal immunity to helminth parasites. Nature 529: 226–230

Graham DB & Xavier RJ (2020) Pathway paradigms revealed from the genetics of inflammatory bowel disease. Nature 578: 527–539

Grencis RK & Worthington JJ (2016) Tuft Cells: A New Flavor in Innate Epithelial Immunity. Trends Parasitol 32: 583–585

Haber AL, Biton M, Rogel N, Herbst RH, Shekhar K, Smillie C, Burgin G, Delorey TM, Howitt MR, Katz Y, et al (2017) A single-cell survey of the small intestinal epithelium. Nature 551: 333–339

Hayakawa Y, Sakitani K, Konishi M, Asfaha S, Niikura R, Tomita H, Renz BW, Tailor Y, Macchini M, Middelhoff M, et al (2017) Nerve Growth Factor Promotes Gastric Tumorigenesis through Aberrant Cholinergic Signaling. Cancer Cell 31: 21–34

Howitt MR, Lavoie S, Michaud M, Blum AM, Tran S V., Weinstock J V., Gallini CA, Redding K, Margolskee RF, Osborne LC, et al (2016) Tuft cells, taste-chemosensory cells, orchestrate parasite type 2 immunity in the gut. Science (80-) 351: 1329–1333

Jiang H, Lei R, Ding SW & Zhu S (2014) Skewer: A fast and accurate adapter trimmer for next-generation sequencing paired-end reads. BMC Bioinformatics 15: 1–12

Kim E, Ahn H, Kim MG, Lee H & Kim S (2017a) The Expanding Significance of Inositol Polyphosphate Multikinase as a Signaling Hub. Mol Cells 40: 315–321

Kim E, Beon J, Lee S, Park SJ, Ahn H, Kim MG, Park JE, Kim W, Yuk J-M, Kang S-J, et al (2017b) Inositol polyphosphate multikinase promotes Toll-like receptor–induced inflammation by stabilizing TRAF6. Sci Adv 3

Kjærgaard S, Jensen TSR, Feddersen UR, Bindslev N, Grunddal K V., Poulsen SS, Rasmussen HB, Budtz-Jørgensen E & Berner-Hansen M (2021) Decreased number of colonic tuft cells in quiescent ulcerative colitis patients. Eur J Gastroenterol Hepatol 25: 817–824

Lee B, Park SJ, Hong S, Kim K & Kim S (2021) Inositol polyphosphate multikinase signaling: Multifaceted functions in health and disease. Mol Cells 44: 187–194

Lei W, Ren W, Ohmoto M, Urban JF, Matsumoto I, Margolskee RF & Jiang P (2018) Activation of intestinal tuft cell-expressed sucnr1 triggers type 2 immunity in the mouse small intestine. Proc Natl Acad Sci U S A 115: 5552–5557

Li B & Dewey CN (2011) RSEM: Accurate transcript quantification from RNA-seq data with or without a reference genome. BMC Bioinformatics

Love MI, Huber W & Anders S (2014) Moderated estimation of fold change and dispersion for RNA-seq data with DESeq2. Genome Biol 15: 1–21

Ma GW, Chu YK, Yang H, Yan XH, Rong EG, L. H & Wang N (2020) Functional Analysis of Sheep POU2F3 Isoforms. Biochem Genet 58: 335–347

Maag D, Maxwell MJ, Hardesty DA, Boucher KL, Choudhari N, Hanno AG, Ma JF, Snowman AS, Pietropaoli JW, Xu R, et al (2011) Inositol polyphosphate multikinase is a physiologic PI3-kinase that activates Akt/PKB. Proc Natl Acad Sci U S A 108: 1391–1396

May R, Qu D, Weygant N, Chandrakesan P, Ali N, Lightfoot SA, Li L, Sureban SM & Houchen CW (2014) Brief report: Dclk1 deletion in tuft cells results in impaired epithelial repair after radiation injury. Stem Cells 32: 822–827

Middelhoff M, Westphalen CB, Hayakawa Y, Yan KS, Gershon MD, Wang TC & Quante M (2017) Dclk1-expressing tuft cells: Critical modulators of the intestinal niche? Am J Physiol - Gastrointest Liver Physiol 313: G285–G299

Nadeem MS, Kumar V, Al-Abbasi FA, Kamal MA & Anwar F (2020) Risk of colorectal cancer in inflammatory bowel diseases. Semin Cancer Biol 64: 51–60

Nadjsombati MS, McGinty JW, Lyons-Cohen MR, Jaffe JB, DiPeso L, Schneider C, Miller CN, Pollack JL, Nagana Gowda GA, Fontana MF, et al (2018) Detection of Succinate by Intestinal Tuft Cells Triggers a Type 2 Innate Immune Circuit. Immunity 49: 33-41.e7

Nagahama Y, Shimoda M, Mao G, Singh SK, Kozakai Y, Sun X, Motooka D, Nakamura S, Tanaka H, Satoh T, et al (2018) Regnase-1 controls colon epithelial regeneration via regulation of mTOR and purine metabolism. Proc Natl Acad Sci U S A 115: 11036–11041

O’Donnell S, Borowski K, Espin-Garcia O, Milgrom R, Kabakchiev B, Stempak J, Panikkath D, Eksteen B, Xu W, Steinhart AH, et al (2019) The Unsolved Link of Genetic Markers and Crohn’s Disease Progression: A North American Cohort Experience. Inflamm Bowel Dis 25: 1541–1549

Qu D, Weygant N, May R, Chandrakesan P, Madhoun M, Ali N, Sureban SM, An G, Schlosser MJ & Houchen CW (2015) Ablation of Doublecortin-Like Kinase 1 in the Colonic Epithelium Exacerbates Dextran Sulfate Sodium-Induced Colitis. PLoS One 10: 1–14

Saiardi A, Erdjument-Bromage H, Snowman AM, Tempst P & Snyder SH (1999) Synthesis of diphosphoinositol pentakisphosphate by a newly identified family of higher inositol polyphosphate kinases. Curr Biol 9: 1323–1326

Sei Y, Zhao X, Forbes J, Szymczak S, Li Q, Trivedi A, Voellinger M, Joy G, Feng J, Whatley M, et al (2015) A Hereditary Form of Small Intestinal Carcinoid Associated With a Germline Mutation in Inositol Polyphosphate Multikinase. Gastroenterology 149: 67–78

Shao L, Oshima S, Duong B, Advincula R, Barrera J, Malynn BA & Ma A (2013) A20 Restricts Wnt Signaling in Intestinal Epithelial Cells and Suppresses Colon Carcinogenesis. PLoS One 8: 1–7

Tesfaigzi J & Carlson DM (1999) Expression, Regulation, and Function of the SPR Family of Proteins: A Review. Cell Biochem Biophys 30: 243–265

Yamashita J, Ohmoto M, Yamaguchi T, Matsumoto I & Hirota J (2017) Skn-1a/Pou2f3 functions as a master regulator to generate Trpm5-expressing chemosensory cells in mice. PLoS One 12: 1–14

Yokoyama JS, Wang Y, Schork AJ, Thompson WK, Karch CM, Cruchaga C, McEvoy LK, Witoelar A, Chen CH, Holland D, et al (2016) Association between genetic traits for immune-mediated diseases and Alzheimer disease. JAMA Neurol 73: 691–697

